# Introduction and validation of a new semi-automated method to determine sympathetic fiber density in target tissues

**DOI:** 10.1101/488338

**Authors:** Dennis Bleck, Li Ma, Lkham-Erdene Byambadoo, Ralph Brinks, Matthias Schneider, Li Tian, Georg Pongratz

**Affiliations:** Hiller Research Center Rheumatology at University Hospital Düsseldorf, Medical Faculty, Heinrich-Heine-University, Düsseldorf, Germany; Neuroscience Center, HiLIFE, University of Helsinki, Helsinki, Finland; Institute of Biomedicine and Translational Medicine, Department of Physiology, Faculty of Medicine, University of Tartu, Tartu, Estonia

## Abstract

In recent years, the role of sympathetic nervous fibers in chronic inflammation has become increasingly evident. At the onset of inflammation, sympathetic activity is increased in the affected tissue. However, sympathetic fibers are largely absent from chronically inflamed tissue. Apparently, there is a very dynamic relationship between sympathetic innervation and the immune system in areas of inflammation, and hence a rapid and easy method for quantification of nerve fiber density of target organs is of great value to answer potential research questions. Currently, nervous fiber densities are either determined by tedious manual counting, which is not suitable for high throughput approaches, or by expensive automated processes relying on specialized software and high-end microscopy equipment. Usually, tyrosine hydroxylase (TH) is used as the marker for sympathetic fibers. In order to overcome the current quantification bottleneck with a cost-efficient alternative, an automated process was established and compared to the classic manual approach of counting TH-positive sympathetic fibers. Since TH is not exclusively expressed on sympathetic fibers, but also in a number of catecholamine-producing cells, a prerequisite for automated determination of fiber densities is to reliably distinct between cells and fibers. Therefore, an additional staining using peripherin exclusively expressed in nervous fibers as a secondary marker was established. Using this novel approach, we studied the spleens from a syndecan-3 knockout (SDC3KO) mouse line, and demonstrated equal results on SNS fiber density for both manual and automated counts (Manual counts: wt: 22.57 +/− 11.72 fibers per mm^2^; ko: 31.95 +/− 18.85 fibers per mm^2^; p = 0.0498; Automated counts: wt: 31.6 +/− 18.98 fibers per mm^2^; ko: 45.49 +/− 19.65 fibers per mm^2^; p = 0.01868). In conclusion, this new and simple method can be used as a high-throughput approach to reliably and quickly quantify SNS nerve fiber density in target tissues.

## Introduction

In order to provide less time-consuming alternatives to the tedious process of manually counting nervous fibers in tissues of interest and to stream line quantification and characterization of nervous fibers, automated and semi-automated processes have been developed and deployed as early as 1979. These processes require special equipment, such as array processors or specialized graphics ports and software, which is highly cost-intensive and often adapted to only one particular purpose (1,2). To allow a more cost-efficient analysis of overall innervation in several target tissues, a semi-automated counting method was established by us. It is based upon several macros programmed for Image J using a basic fluorescence microscopy set up.

Usually, tyrosine hydroxylase (TH) is used as a marker of sympathetic fibers. TH catalyzes the conversion from L-tyrosine to L-3,4-dihydroxyphenylalanine (L-DOPA), which represents the rate-limiting step of catecholamine synthesis (3,4). TH is ubiquitously expressed in sympathetic nervous fibers as well as in a multitude of other cells throughout most mammalian tissues. Due to the fact that TH is not exclusively expressed on nervous fibers, we decided to introduce a counterstaining to improve distinction between TH-positive sympathetic fibers and TH-positive cells. As a neurofilament, ubiquitously and exclusively expressed in nervous fibers, peripherin is an excellent candidate for double stainings (5). We hypozethized, that with peripherin and TH co-staining, TH-positive fibers will be distinguished from TH-positive cells, since fibers will be discernible by the co-localization of peripherin and TH, while all other TH signals originate only from TH-positive cells. If high-end technology required for automated counting processes is not available, fiber density is usually determined by manually counting visible TH-positive fibers in 17 high power fields (HPF), according to published methodology (6). However, this is a time-consuming and observer-dependent process. We present in this work a simple, high-throughput, automated screening method of sympathetic fiber density in tissues.

The nervous system plays a major role in regulating immune responses. Since the early 1980s, the sympathetic nervous system (SNS) innervation of lymphoid tissues has been investigated, particularly in rats (7–10). Sympathetic fibers were discovered in the vasculature and in the parenchyma in close proximity to lymphoid effector cells within the primary and secondary lymphoid organs(8). The neuroimmune junction between SNS fibers and immune cells has been described as about 6 nm in width, which strongly suggests a direct effect of SNS neurotransmitters on cells of the immune system (8). The neurotransmitters of the SNS are epinephrine (E) and norepinephrine (NE), also known as adrenalin and noradrenalin. The precursor of both molecules is dopamine and all three molecules are derived from the amino acid L-tyrosine and are summarized as catecholamines. Catecholamines are ligands of adrenoceptors. Between E and NE, the latter plays the major role as neurotransmitter of the SNS. Extensive research has been conducted to prove that NE facilitates neurotransmission from the SNS to immune cells (11). Functional adrenergic receptors are discovered on cells of both the innate and the adaptive immune systems (12). Besides local secretion from SNS fibers, high amounts of E and NE are synthesized by chromaffin cells in the medulla of the adrenal gland and released into the circulation. The adrenal gland is the final effector of the so-called hypothalamic-pituitary-adrenocortical (HPA) axis. This circuit represents the other major central nervous system (CNS)-controlled pathway to regulate immune functions, next to the SNS (13).

Next to important functions such as degradation of senescent erythrocyte and subsequent iron recycling, the spleen is also involved in key processes for the development, maturation and homeostasis of the immune system. Within the follicles of the white pulp of the spleen, germinal centers are formed. These globular structures are the site of B-cell maturation and more importantly antibody isotype switching and antigen affinity refinement(14). SNS fibers reach the lymphoid parenchyma of the spleen after branching off the neurovascular plexuses along the local vasculature in the tissue. Noradrenergic fibers are found in the periarteriolar lymphatic sheath (PALS) surrounding the central artery. This is a T-cell rich area, in which germinal centers develop to very dynamic structures. SNS fibers can be found in the marginal zones but not inside of the germinal centers (13,15). This could be explained by the high level of proliferation taking place in the germinal centers. A rapid expansion of B-cells potentially forces the nervous fibers to the edges of the germinal centers.

The degree of sympathetic innervation is very dynamic as is evidenced by the fact that, during acute local inflammation, the fiber density is decreased in the affected tissue, while systemic sympathetic activity is increased (16). Upregulation of the SNS can lead to cardiovascular hypertension and heart failure (17,18). Alterations in the degree of sympathetic innervation have also been described in chronic kidney disease and in the spleens of acute sepsis patients (19,20). Little is known about the mechanisms regulating the degree of sympathetic innervation, however. Syndecan-3 (SDC3), also known as N-Syndecan, is a member of a family of transmembrane heparan sulfate proteoglycans, a group of cell surface molecules mostly responsible for cell – extracellular matrix contact and interaction. They are closely related to heparin, which is known for its clinical use as an anticoagulant, due to its capacity to bind to a large number of proteins, such as chemoattractant growth factors and cytokines (21). SDC3 is involved in the cortactin-Src kinase-dependent and epidermal growth factor receptor-induced axonal outgrowth and cell migration during development of the brain (22–24). Furthermore, Sdc3-knockout (SDC3KO) mice have been shown to be resistant to diet-induced obesity (25-27)and cocaine-abuse(28), all of which are tightly regulated by the autonomic nervous system. We hence hypothesize that SDC3 might play a role in regulating the autonomic nervous activity and in particular the sympathetic innervation in target organs.

## Materials and Methods

### Animals

C57B/6J littermate mice were used as wild type control. SDC3KO mice (C57BL/6J) were generated at the Neuroscience Center, University of Helsinki, Helsinki, Finland under approval by the National Animal Experiment Board of Finland under the license number ESAVI/7548/04.10.07/2013 and ESAVI/706/04.10.07/2015, and the methods were carried out in accordance with the approved guidelines.

### Tissue preparation and cryo sections

Mice were sacrificed by CO_2_ and target organs and tissues were immediately isolated from the animals. The tissues were placed in specimen molds and covered in embedding media. The molds were then placed in liquid nitrogen for shock freezing. For sections containing bone marrow, the femurs were isolated and put into 4% paraformaldehyde (PFA) for 48 hours at +4°C. Then specimens were washed with dest H_2_O and transferred into 15% EDTA for decalcification for 48 hours at +4°C. After washing again with dest H_2_O, they were placed in a 25% sucrose solution overnight until they had sunken to the bottom of the tube. For preparation of cry sections, specimens were removed from molds and mounted onto the specimen disk of the cryostat using embedding medium. After the embedding medium was frozen hard, trimming was started at 50 μm until the tissue was visible. The trimming steps were then gradually reduced to 30 μm, 15 μm, 10 μm and 5 μm. Sections were then produced at 3 μm. Sections were raised onto microscope slides and then placed on top of the cryostat and kept there for about 10 minutes to dry. Afterwards tissues were immediately fixated by placing the tissue sections in paraformaldehyde (PFA) 3% for 15 minutes at RT. Then, the slides were washed in PBS (1×) for 10 minutes at RT.

### Immunofluorescence staining

Sections were stained using rabbit polyclonal anti TH ab152 (Merck, Darmstadt, Germany) and chicken polyclonal anti peripherin ab39374 (Abcam, Cambridge, UK) antibodies over night after blocking with 2 % normal goat serum and 0.3% Triton X in PBS. Secondary antibodies goat anti rabbit alexa fluor 594 (Invitrogen, Darmstadt, Germany) and goat anti chicken IgY alexa fluor 488 (Abcam, Cambridge, UK) were used for labeling and cover slips were mounted using ProlongGold containing DAPI (Invitrogen, Darmstadt, Germany).

Zeiss Axioscop 2 plus (Carl Zeiss AG, Oberkochen, Germany) with Nikon DSV VI1 camera and Nikon imaging system (NIS) freeware software (Nikon, Düsseldorf, Germany) were used for documentation. Images were taken 24 hours after mounting of the cover slips. Negative controls and isotype control stainings were analyzed first for each staining and each tissue. The duration of exposure was set according to these controls and all samples of one tissue were recorded with the same setting. Images were saved as lossless tagged image format files (.tif). For this project all merged images were created using the open Image J software (29,30). Brightness and contrast settings were augmented to the same values for all images shown in this document. Scale bars were set using Image J and panels were generated with Adobe Illustrator CS2.

### Data collection and statistical analysis

TH positive fibers and cells were counted using two different approaches. The classical approach was to manually count fibers in 17 random HPF according to published protocols (6). In order to develop automated counting processes for fibers and cells, respectively, algorithms were created using the Image J platform (see supplement 1 for program code; figures S1-S4). All results were analyzed using R statistical software package, version 3.5.0 (The R Foundation for Statistical Computing). Statistical significance was determined by two-sided t - tests and p-values below 0.05 were considered significant.

## Results

### Sympathetic fiber counts are higher with a semi – automated method as compared to a manual process, and simultaneous TH-positive cell counting is possible

Figure 1 shows an example of TH – peripherin co – staining in wt spleen sections. The co – staining allows for identification of double positive sympathetic fibers (indicated by yellow arrows). At the same time, it provides the opportunity to differentiate fibers from TH-positive cells in the tissue, since those cells are peripherin-negative (indicated by red arrows). Examples of images from all other tissues included in this study can be found in the supplement (Figures S5 – S9).

**Figure 1:**
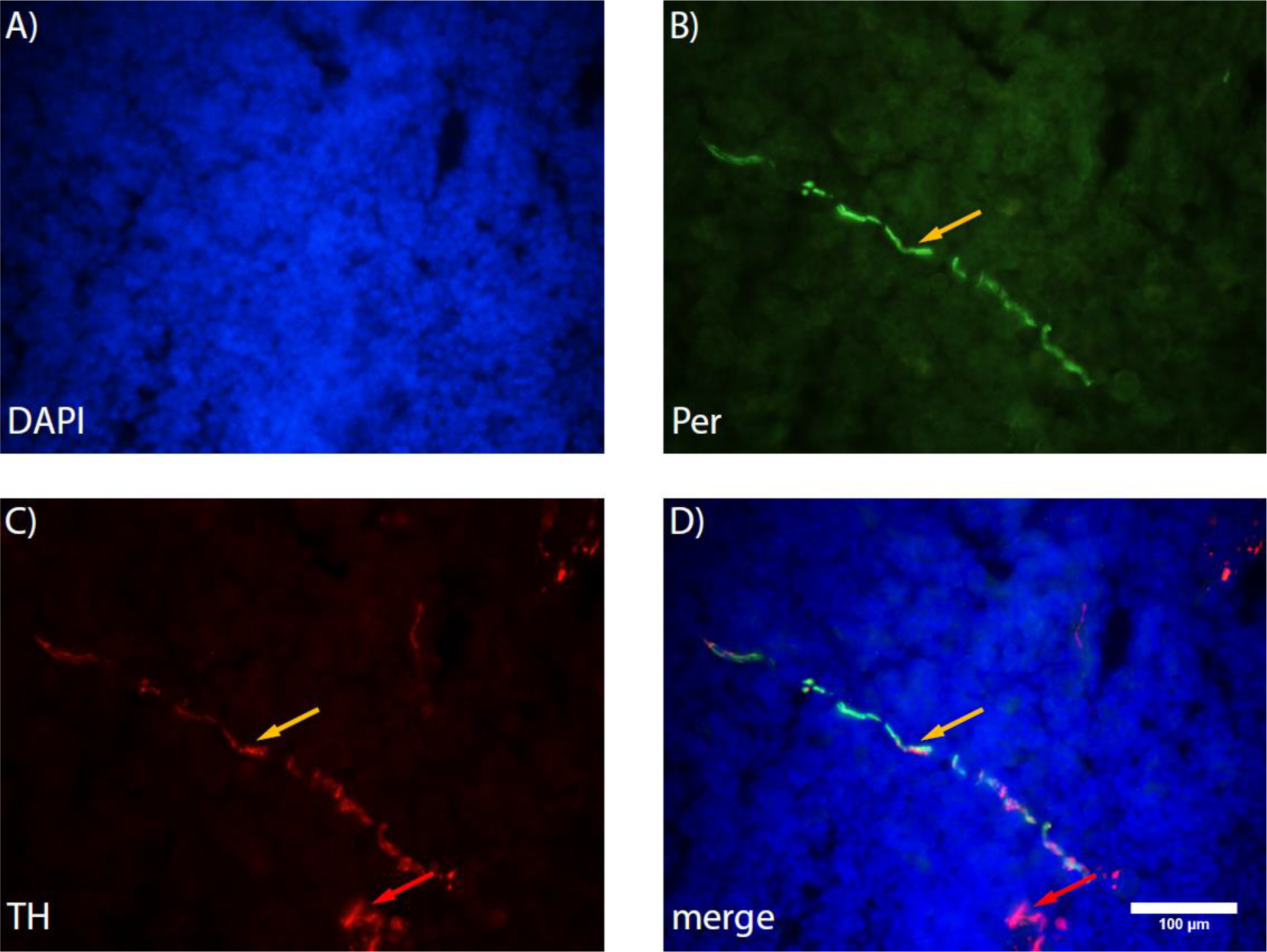
Exemplary image of TH – peripherin co – staining in a spleen section. A) Nuclei are labeled with DAPI. B) Peripherin is labeled green (alexa fluor 488). C) TH is labeled red (alexa fuor 594). D) Merged image. Magnification is 400 fold. Red arrows indicate TH positive cells, Yellow arrows indicate sympathetic fibers.

Fibers were counted in 24 images of every tissue from wild type animals. Either by eye (manual counts, mc) or using Image J (automated counts, ac), fibers were defined as objects of oblong shape and at least 50 μm in length only for mc. Each image has a size of 1600 pixels in width and 1200 pixels in height. Due to the scale of 2.828 pixels per μm at a magnification of 400 ×, each image covers a slice of 0.24 mm^2^. Overall, the automated method shows a higher sensitivity and acquires significantly more fibers than the manual approach (Figure 2, A), p = 0.009772). This is due to the higher counts in the bone marrow and thymus (figure 2, C) and G), p = 0.00034 and p = 0.04481). The TH – peripherin co – staining allows not only for determination of fiber densities, but also offers an opportunity to count TH-positive cells. As shown in figure 2H, the tissue with the most TH-positive cells per image was the submandibular gland, with an average of 313.02 +/− 83,76 cells per mm^2^, followed by the spleen with an average of 194.97 +/− 93.77 cells per mm^2^. The least amount of TH-positive cells was counted in the heart (94.27 +/− 53.28 cells per mm^2^). In the bone marrow, adrenal gland and thymus sections, 94.62 +/−75.87, 131.25 +/− 67.61 and 96.01 +/− 48.36 cells per mm^2^ were recorded, respectively.

**Figure 2:**
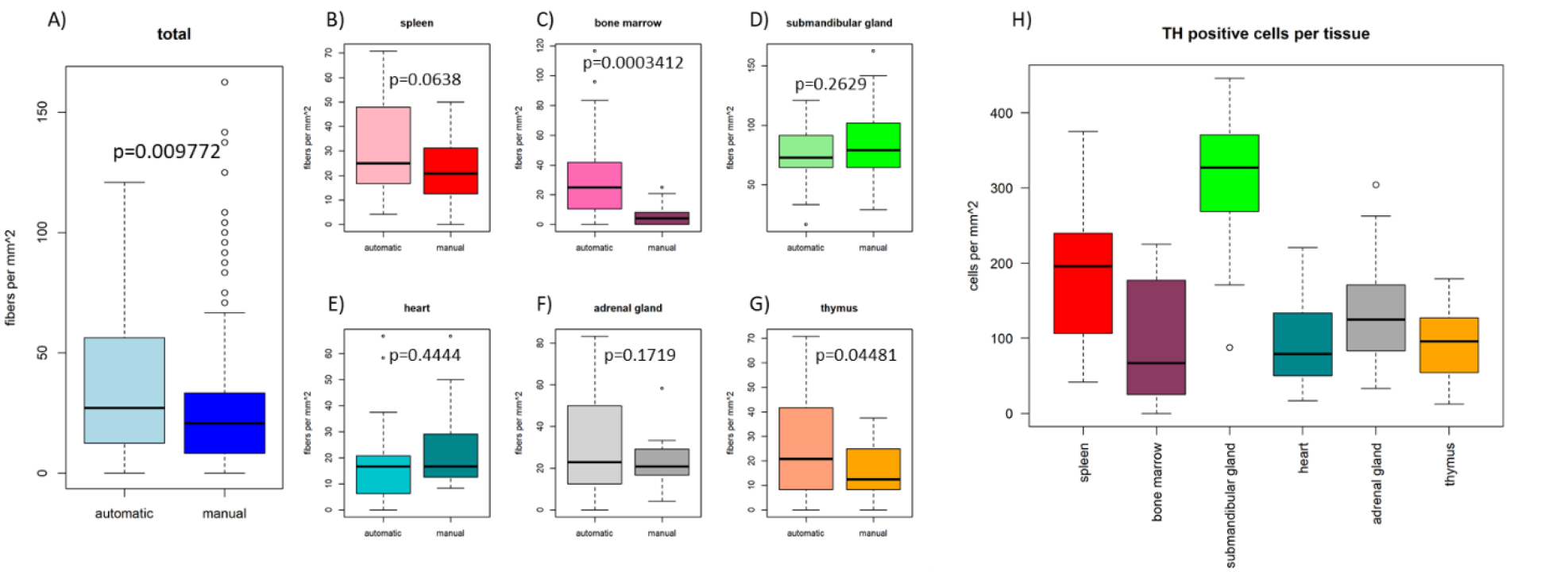
Fiber counts acquired by manual counting and automated approach and TH positive cells in each tissue. A) Shows the total fibers counted with both approaches. The automated approach acquires significantly more fibers than the manual method (p = 0.009772). B) Fiber counts in the spleen. The automated method yields a higher fiber count than the manual approach (p = 0.0638). C) in bone marrow automated counts are significantly higher than manual counts (p = 0.0003412). D) In submandibular gland, there is little difference between both methods (p = 0.2629). E) Similar counts were acquired by both methods in heart sections (p = 0.4444). F) In adrenal gland sections, both methods yield similar fiber counts (p = 0.1719). G) Automated counts are significantly higher than manual counts in thymus sections (p = 0.04481). P - values were determined by two-sided t - test. TH positive cells per mm^2^ in each individual tissue (H)). The highest numbers of cells were counted in images of submandibular gland and spleen sections. The lowest counts were acquired in images of sections from heart.

Plotting the number of manual fiber counts versus the number of automated fiber counts per image shows that overall the automated method picks up more fibers per image than the manual approach (figure 3, A). This is illustrated by the linear regression line with a positive slope of 0.81. Most of the counts above the regression line were acquired in images of the submandibular gland and heart sections.

**Figure 3:**
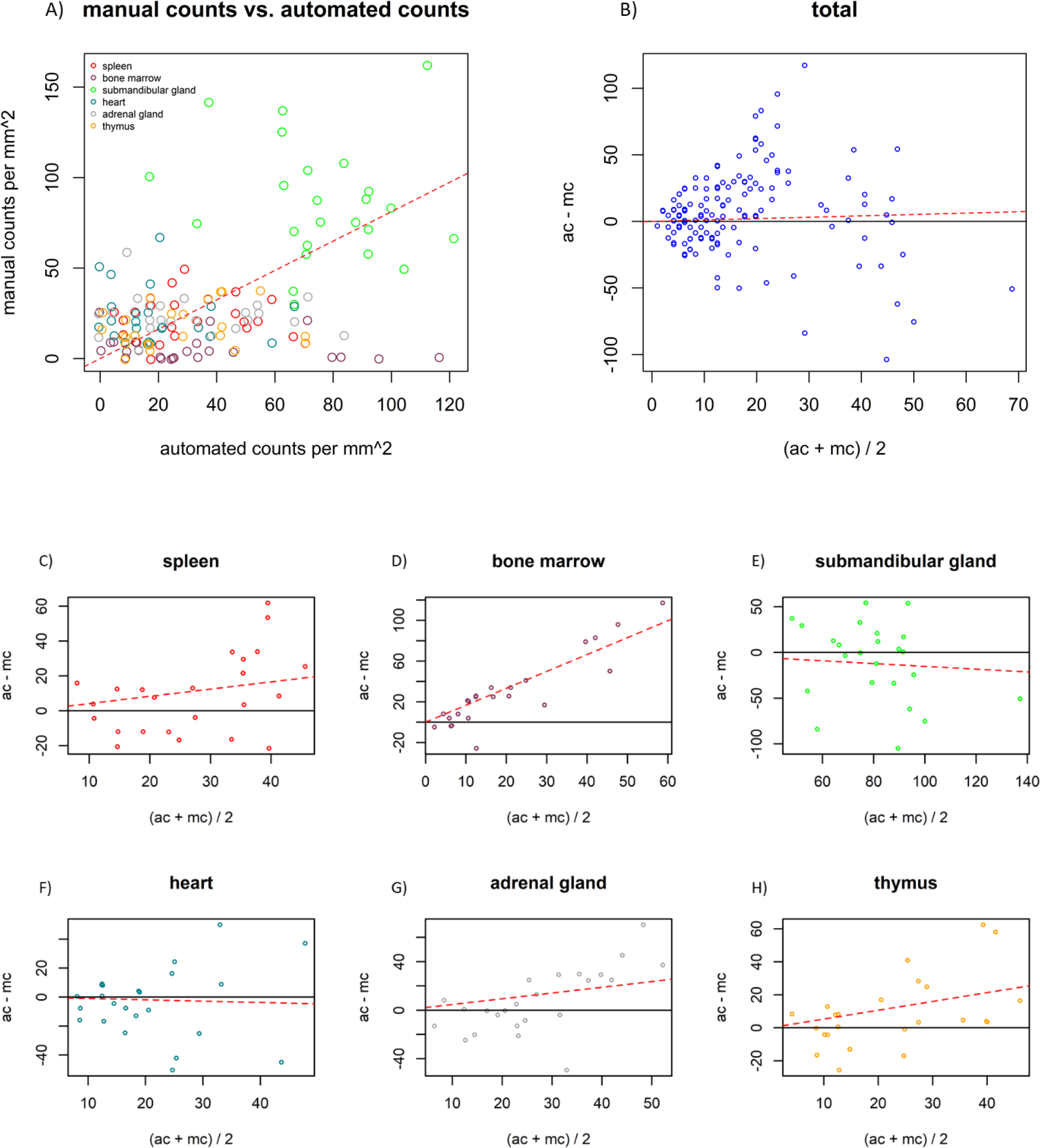
Scatter plot and Bland - Altman plots of manual count values vs. automated count values per image overall and in each individual tissue. A) The regression line of the scatter plot has a slope of 0.81 (red dotted line), illustrating higher fiber counts per image with the automated method compared to the manual approach. B) the Bland - Altman plot shows that automated counts (ac) were higher than manual counts (mc) overall. Slope of the regression line is 0.10 (red dotted line). C) automated counts (ac) are higher than manual counts (mc) in spleen. The slope of the regression line is 0.41 (red dotted line). D) ac are higher than mc in bone marrow. The slope of the regression line is 1.66 (red dotted line). E) ac are lower than mc in submandibular gland. The regression line has a slope of −0.15 (red dotted line). F) In heart sections ac are lower than mc. The regression line has a slope of - 0.09 (red dotted line). G) ac are higher than mc in adrenal gland sections. The regression line has a slope of 0.47 (red dotted line). H) In thymus sections, ac are higher than mc illustrated by a regression line with a slope of 0.53 (red dotted line).

The Bland – Altman plot shows that overall automated counts were higher than manual counts by plotting the difference between ac and mc versus the mean of ac and mc (figure 3B). The slope over all tissues is 0.10. In figure 3C, the slope is 0.41 in the spleen sections. This tendency was similar in the bone marrow, adrenal gland and thymus sections (figure 3, D), G) and H)). The slopes of the regression lines are 1.66, 0.47 and 0.53, respectively. The slope of 1.66 for the bone marrow being the steepest, indicating that the automated method acquires more total events in sections where the average events are increased. The slopes of the regression lines in the Bland – Altman plots were negative for the submandibular gland (−0.15) and heart (−0.09) sections (figure 4, E) and F)), indicating that more events were acquired by the manual method as the average number of events was increased in these sections. Overall, the Bland – Altman plots show that the automated method is slightly superior to the manual method.

**Figure 4:**
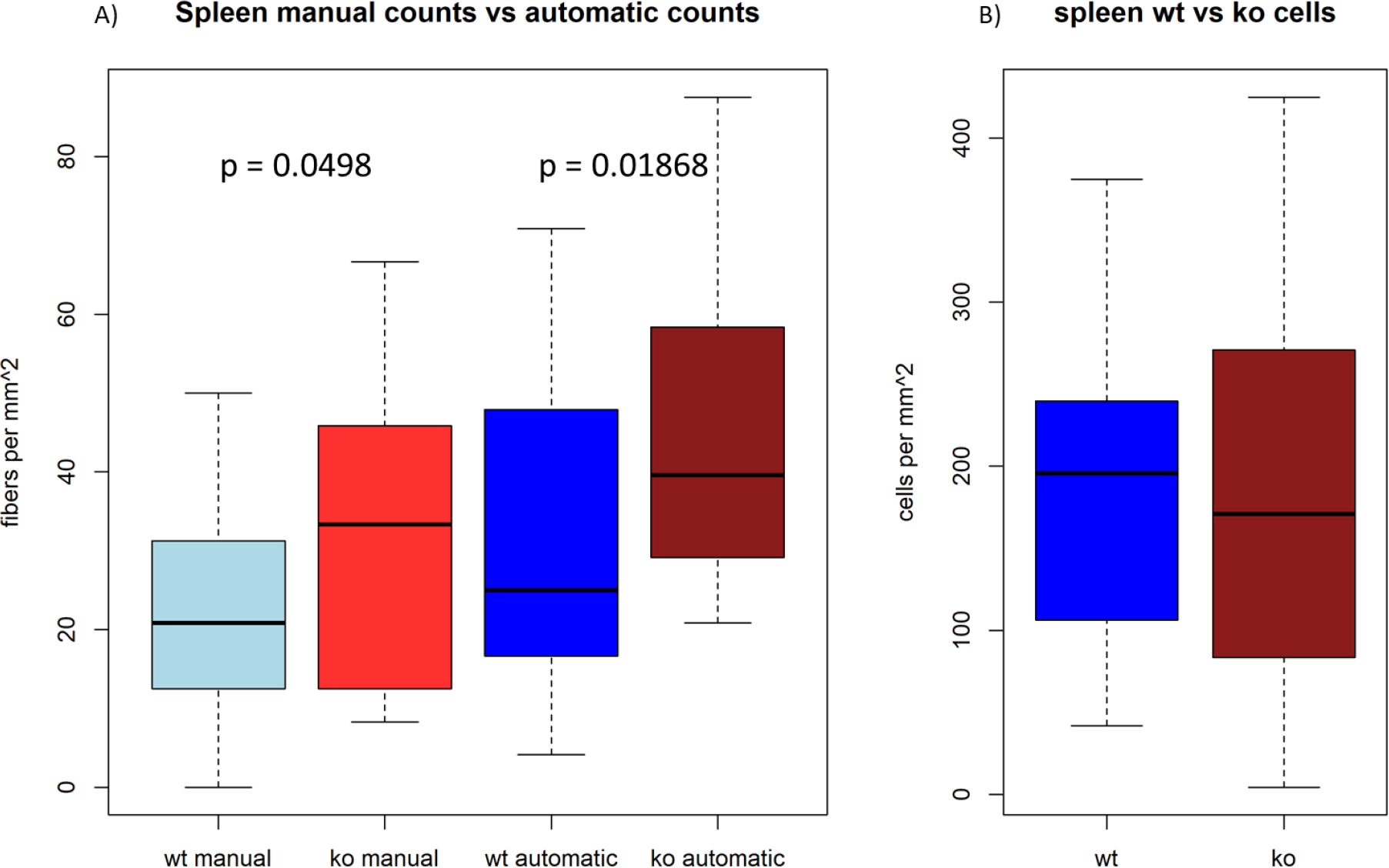
Sympathetic fiber and TH positive cell counts in spleens from wt and SDC3KO mice acquired by manual and automated process. A) Sympathetic fiber density is significantly increased in SDC3KO (red boxplots) compared to wt (blue boxplots). For manual counts p = 0.0498; for automated counts p = 0.01868. Overall, a significantly higher fiber density is recorded by the automated method with a p value of p = 0.001319. B) TH positive cell number is not different in SDC3KO spleens (red boxplot) compared to wt spleens (blue boxplot).

### Sympathetic fiber and TH-positive cell counts are higher in the SDC3KO spleens compared to the wt spleens

As depicted in figure 4A), sympathetic fiber density is significantly increased in the spleens from SDC3KO mice compared to wt mice. This is represented by both counting methods. For the manual approach the p value was p = 0.0498 while the automated procedure resulted in a lower p value of p = 0.01868. Overall, the automated method registered significantly more fibers compared to the manual process with a p value of p = 0.001319. At the same time, TH-positive cell number is not increased in the spleens of SDC3KO mice compared to the spleens of wt mice (figure 4B). These results hint to an increased sympathetic activity in the spleen due to a loss of SDC3.

## Discussion

The automation of the counting process for sympathetic nerve fibers and TH+ cells, respectively, presents a number of advantages. First of all, it is considerably less time-consuming than counting fibers, or other target structures, by eye. It also eliminates the effect of subjective perception by the experimenter from the process. Another benefit of the automated process is the fact, that the images captured for the analysis are available for future studies or replication of the analysis, while in the previously described manual approach, targets were counted under the microscope without capturing the area of investigation as images (6). Compared to other automated processes that have been used to count and analyze nervous fibers in tissue sections, this approach does not require any special equipment or software. It is therefore a lot more cost-efficient than other approaches, which are based on three-dimensional analysis (31–33). Apart from these practical advantages, the automated method offers a number of analytical upsides. With the automated approach, it is possible to count double-positive structures of variable shapes and sizes, whereas the previous manual method only allowed for a discrimination by size and shape, for example by only counting objects that were fiber-shaped and above 50 μm in length determined through a micrometer eyepiece (6). These discrimination criteria eliminate all fibers running perpendicular to the plane of the section, which will be the largest proportion of fibers, and only the least number of fibers running horizontally to the plane of the section is registered for analysis. Therefore, the number of fibers recorded by the automated approach is increased, due to the fact that all double-positive structures were registered.

While the novel method described here has many advantages over the manual approach, a few shortcomings should be mentioned in comparison to other automated methods. While other processes might be more expensive, they do offer opportunities for more detailed analysis than the simple method presented here. For example, a three-dimensional analysis of fibers within the analyzed tissues allows for determination of exact number of fibers and branches or junctions. With methods that are more elaborate, it is possible to observe structural changes, such as fiber diameter, when comparing different groups (treatment, knock out, etc.). In addition, fibers could be tracked to their origin, which opens up opportunities to further analyze fiber functions. In case these additional details are not important for analysis, however, the approach described in this paper offers a simple and fast alternative in order to determine fiber densities in target tissues.

Overall, the automated method acquires significantly more fibers than the manual approach. When considering the results for the individual tissues, this trend is confirmed in the bone marrow and thymus sections. In the spleen, submandibular gland, heart and adrenal gland sections, no significant difference is apparent. The two methods seem to be nearly equal in these tissues. The scatter blot shows that in general, the automated approach counts more fibers per image than the manual approach. This finding is confirmed by the Bland – Altman plots showing that overall the automated approach is slightly superior to the manual approach. When comparing the methods in each of the analyzed tissues individually, the same trend is visible for the spleen, bone marrow, adrenal gland and thymus sections. Only in tissues with extreme degrees of innervation, the submandibular gland and heart, the manual approach acquired more fibers than the automated approach. This is assumedly due to the fact, that manual counts are inaccurate due to the complexity of tissues with such high fiber densities (34–38). Another upside due to the introduction of the double-staining is the possibility to count TH-single positive cells by blanking peripherin-positive areas from the image before analysis. Due to very high cell densities in some tissues, an evaluation by eye is difficult and less reliable.

Both methods showed a significant increase in fiber density in the spleens from SDC3KO animals. This demonstrates that the automated method is at least as reliable as the previously used manual method. The increase in innervation in the spleens of SDC3KO animals suggests surprisingly a growth inhibitory function of SDC3. This is however not supported by previous findings on neurite-promoting function of membrane-bound Sdc3 in the brain. N-syndecan serves as a receptor or a co-receptor for heparin-binding growth associated molecule (HB-GAM, also known as pleiotrophin) (39) and that addition of exogenous heparin, as well as heparitinase treatment of neurons, both inhibit HB-GAM-induced neurite outgrowth (40). Kinnunen et al. from 1996 further showed that exogenous SDC3 isolated from perinatal rat brains inhibited-HB-GAM dependent neurite outgrowth in vitro(23). A significant survival-deficiency of dorsal root ganglion neurons was described for SDC3KO mice during the first postnatal week, whereas neurons isolated from young adult SDC3 KO mice showed no reduction in survival compared to WT controls (24). Since all data are generated in the CNS of young animals the mechanism of enhanced peripheral sympathetic innervation in adult SDC3KO spleens observed in this work is still elusive to us and awaits a further depiction. In contrast to all previous data our findings suggest an inhibitory function of SDC3 in adult peripheral sympathetic tissue.

Overall the introduction of a secondary staining and the software-based analysis results in a time-saving, highly objective and flexible method to count structures of any shape and size in tissue sections. The method is therefore suitable for a wide variety of applications when analyzing SNS innervation and TH+ cells in a variety of tissue sections in health and disease. At the same time, it does not rely on expensive software or microscopy equipment.

## Acknowledgements

We thank Ellen Bleck and Birgit Opgenoorth for excellent technical assistance.

## Supplement 1

### Macros

#### Determination of TH positive fiber density using peripherin and TH double staining

One macro was deployed to select TH positive areas in the images:

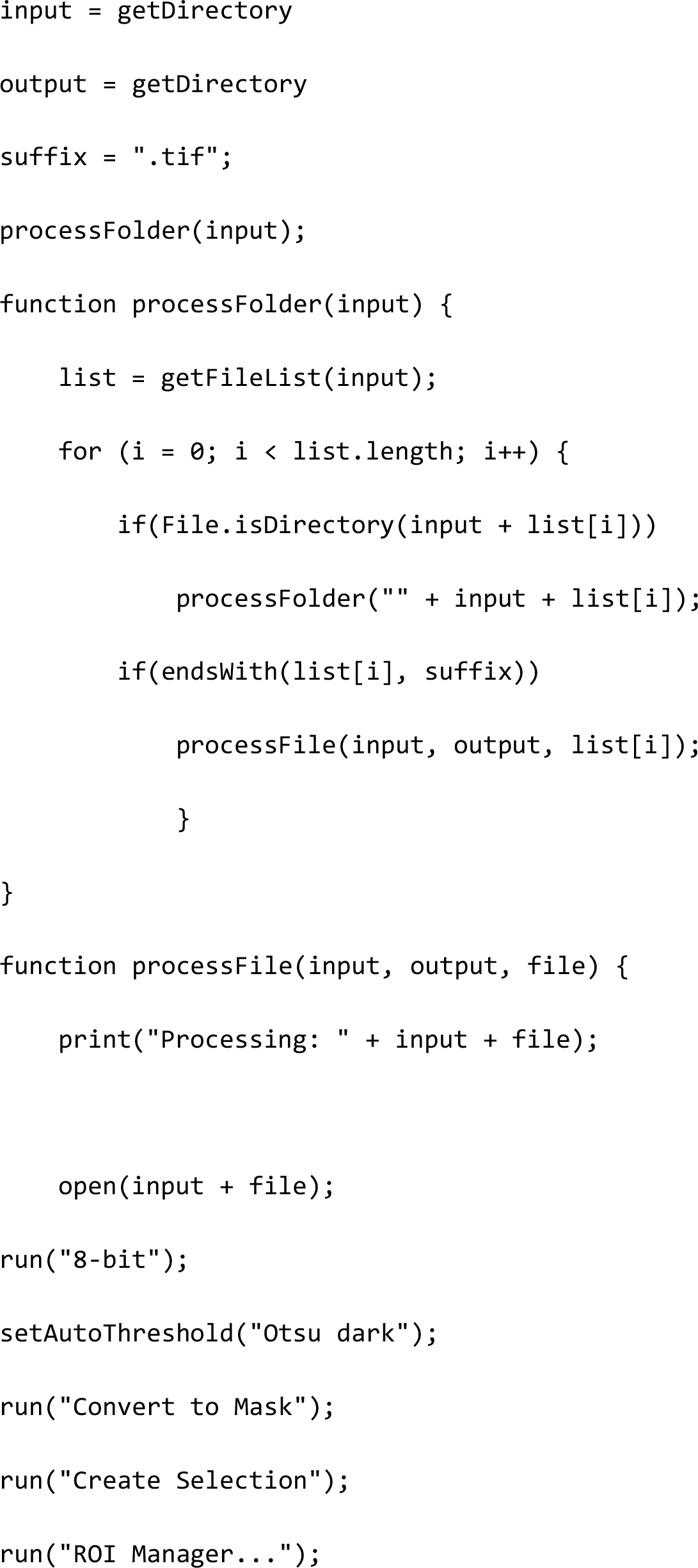

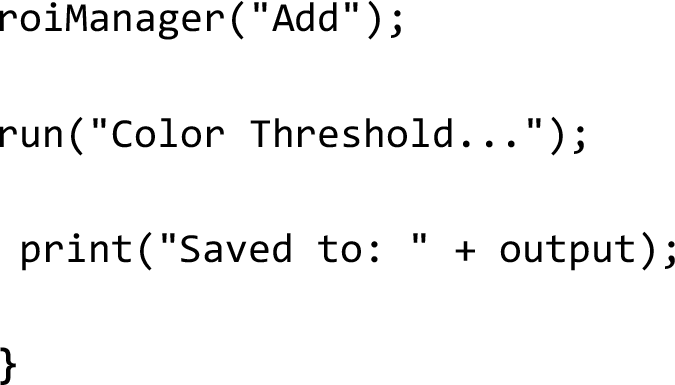

The ROIs created in the first macro were placed over the peripherin image to determine where both stainings were co-localized.

Fiber count:

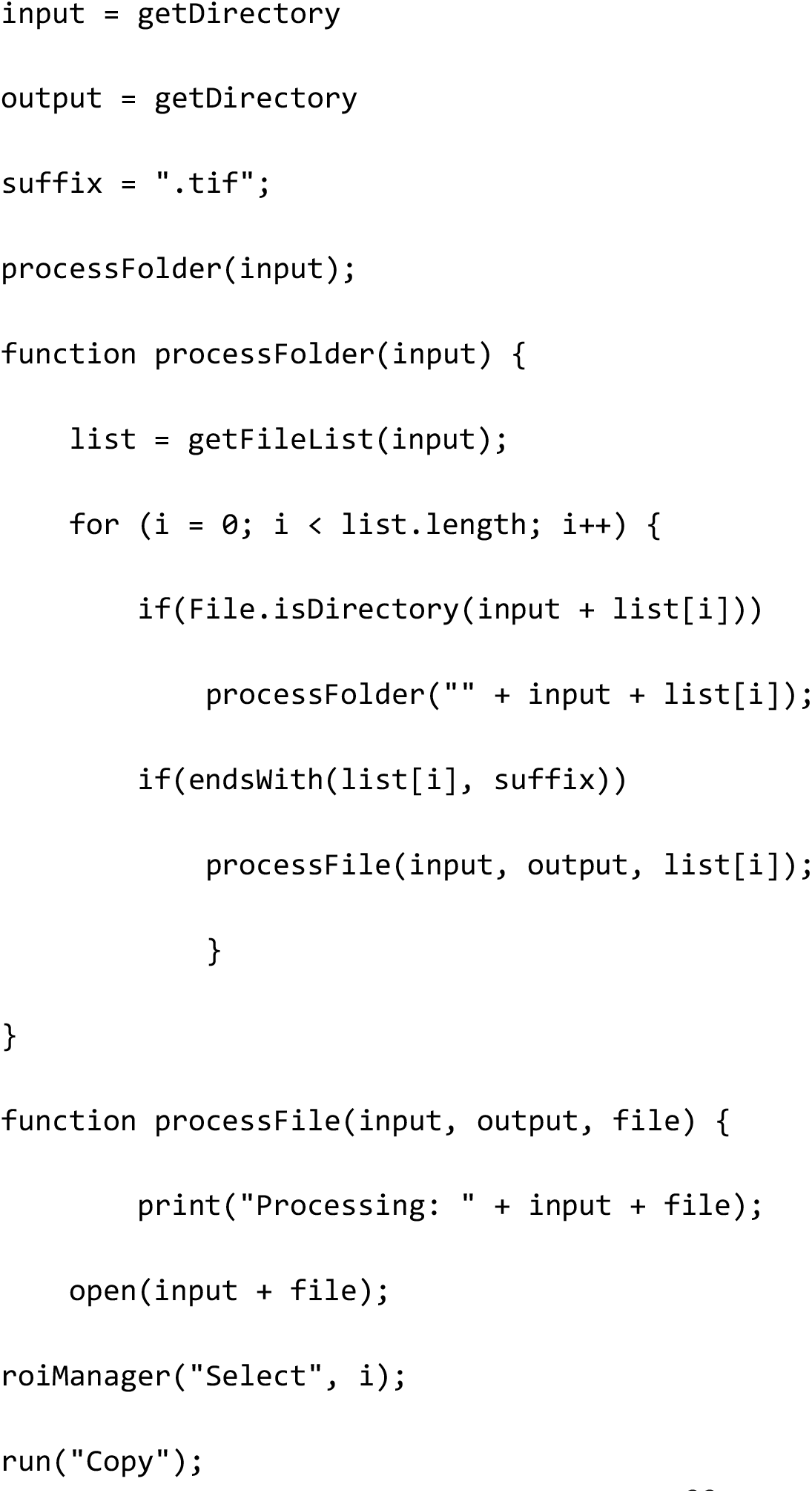

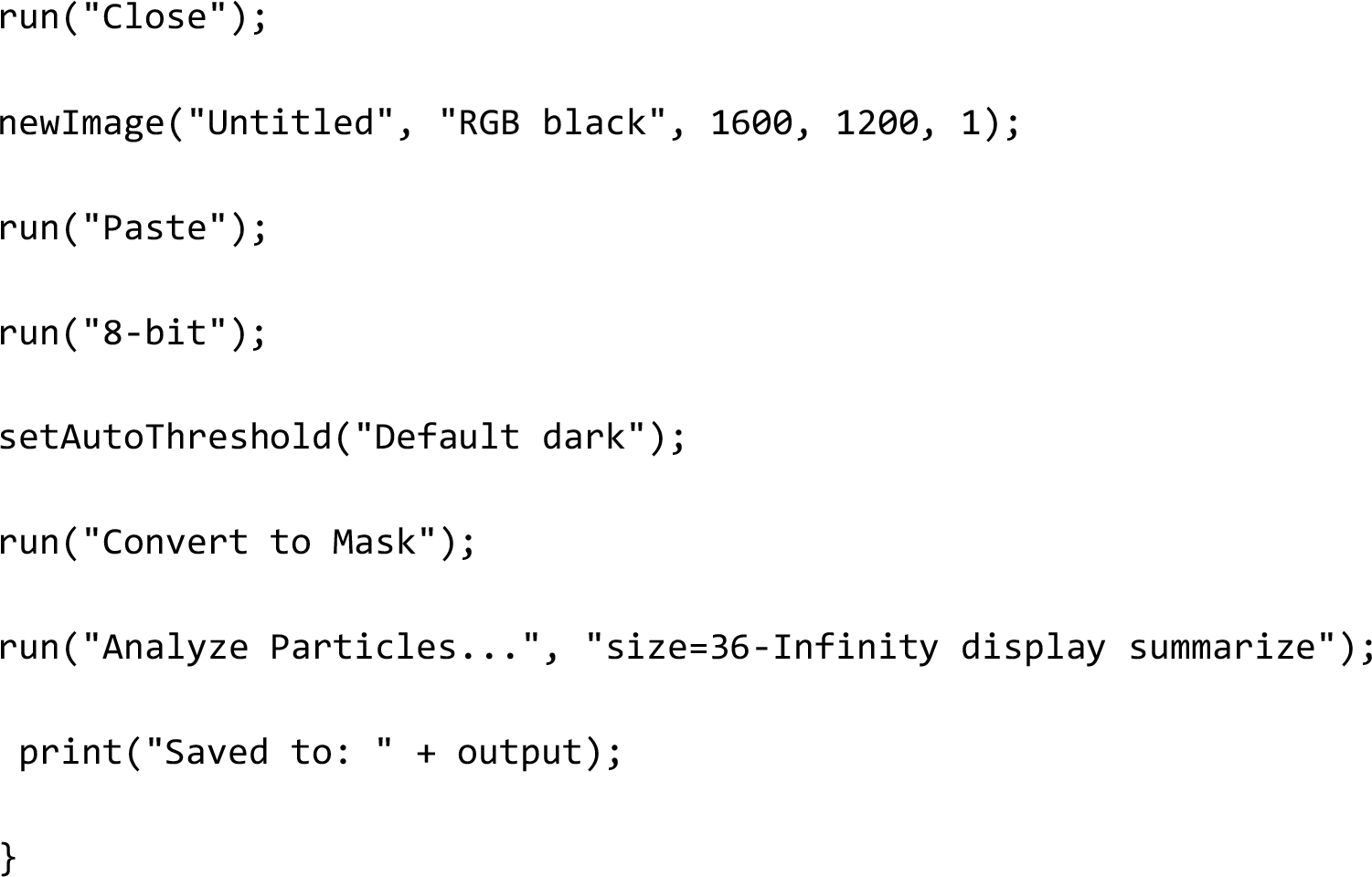

#### Determination of TH positive cell quantity in TH and peripherin double stained sections

First macro was used to select peripherin positive areas:

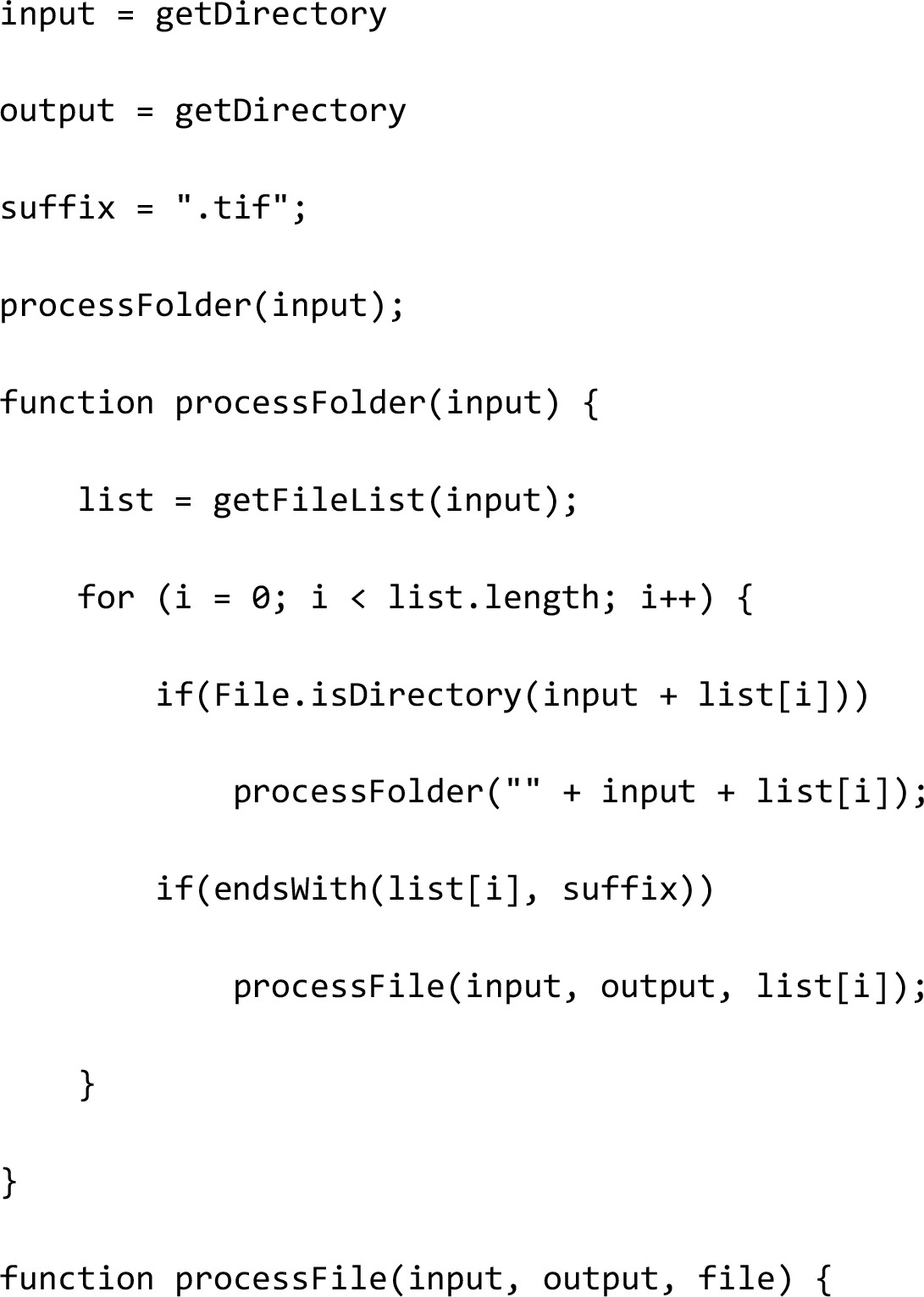

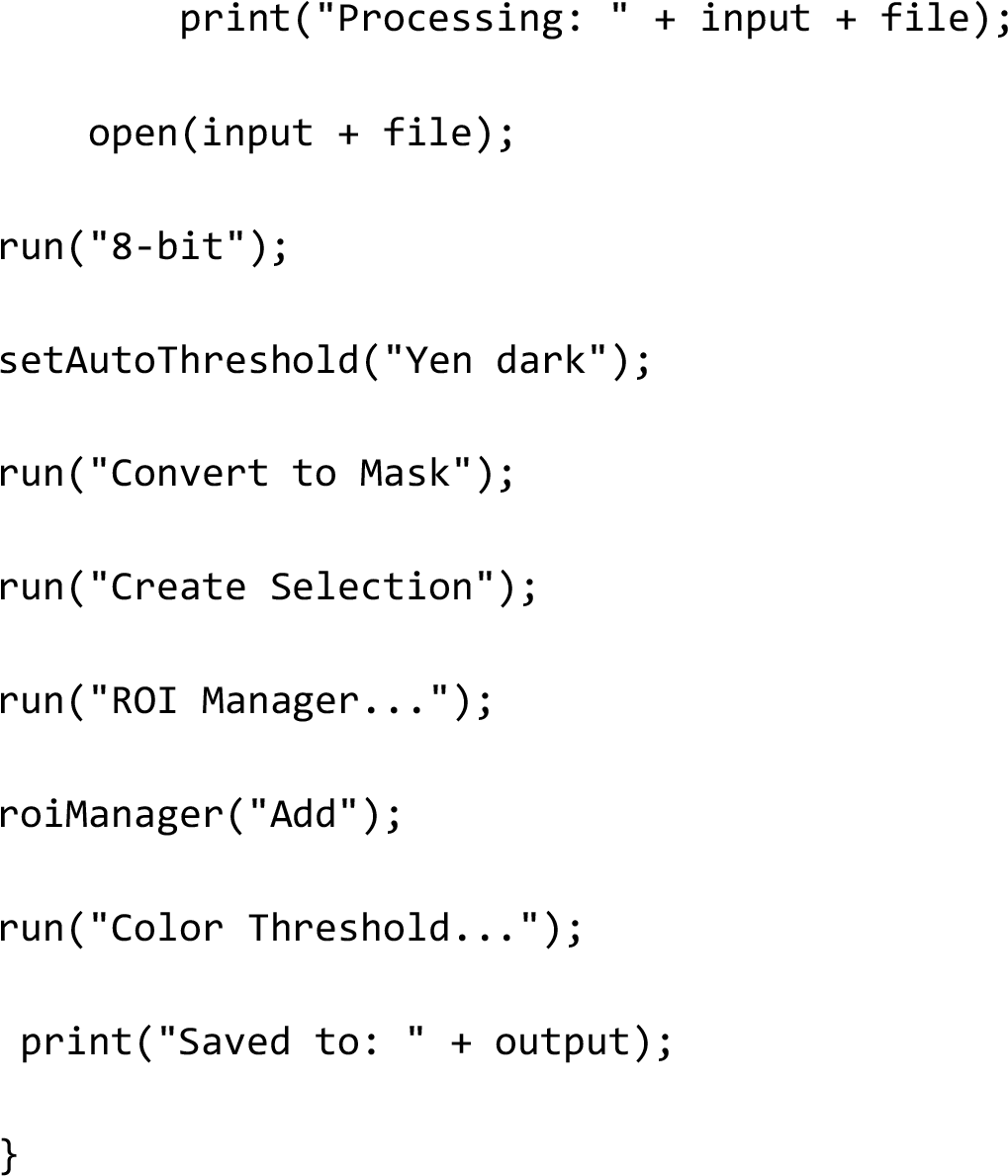

A second macro was used to place the ROIs created from the first algorithm over the TH staining images and clearing these selected areas. All TH positive areas that were left were single positive and therefore considered TH positive cells as opposed to fibers.

Clearance of peripherin positive areas and TH positive cell count:

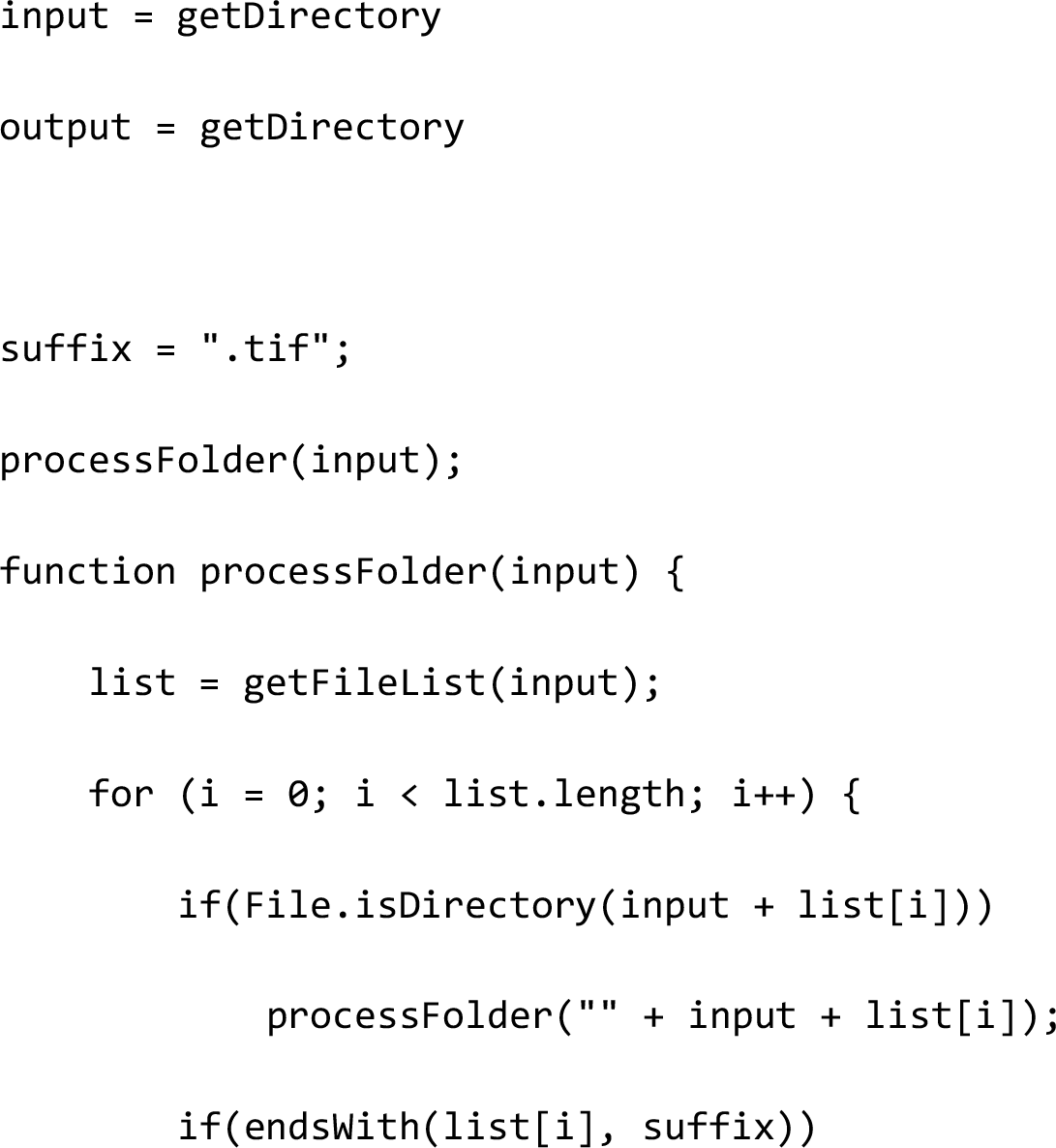

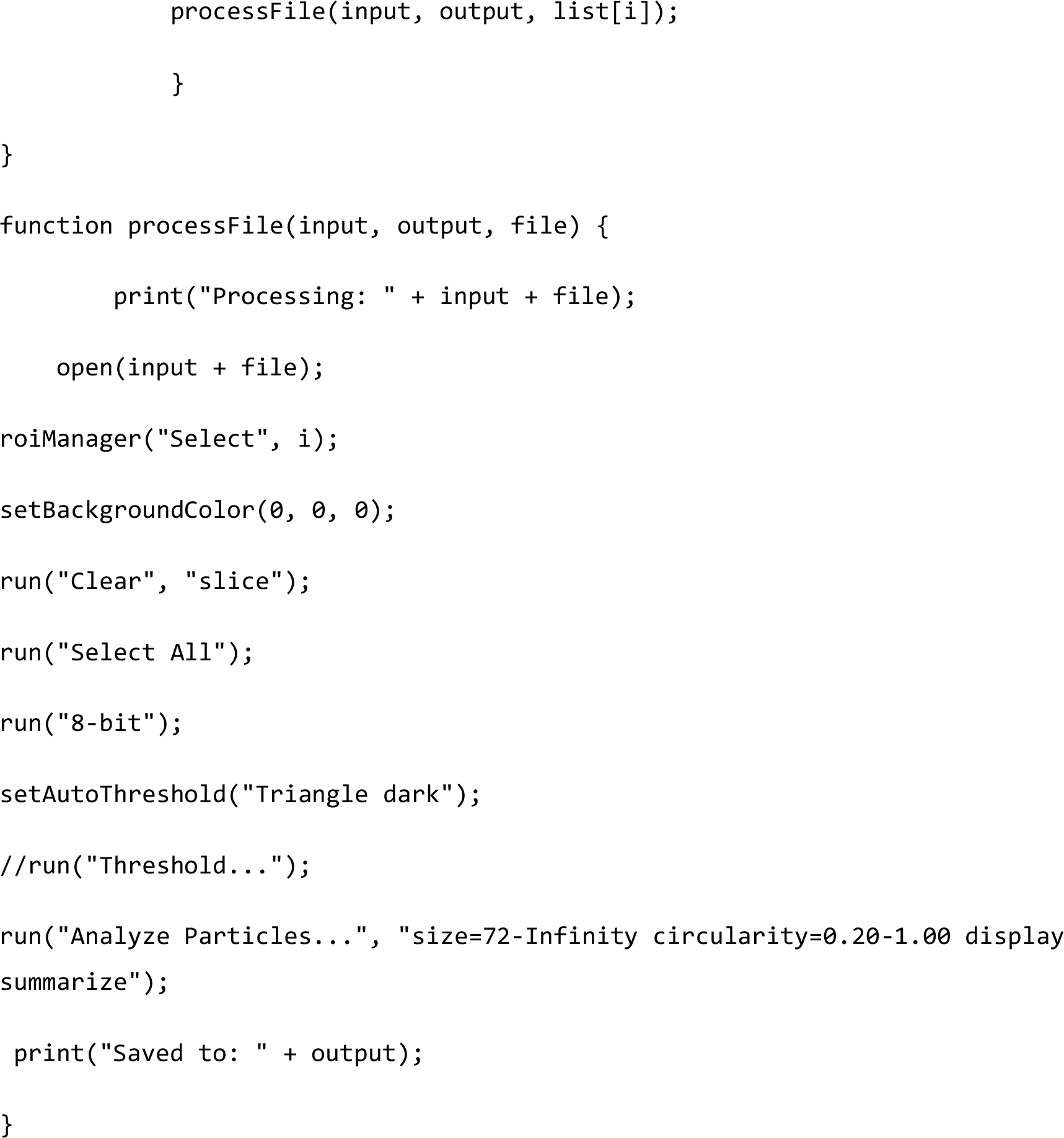

## Supplementary figures

**Figure S1:**
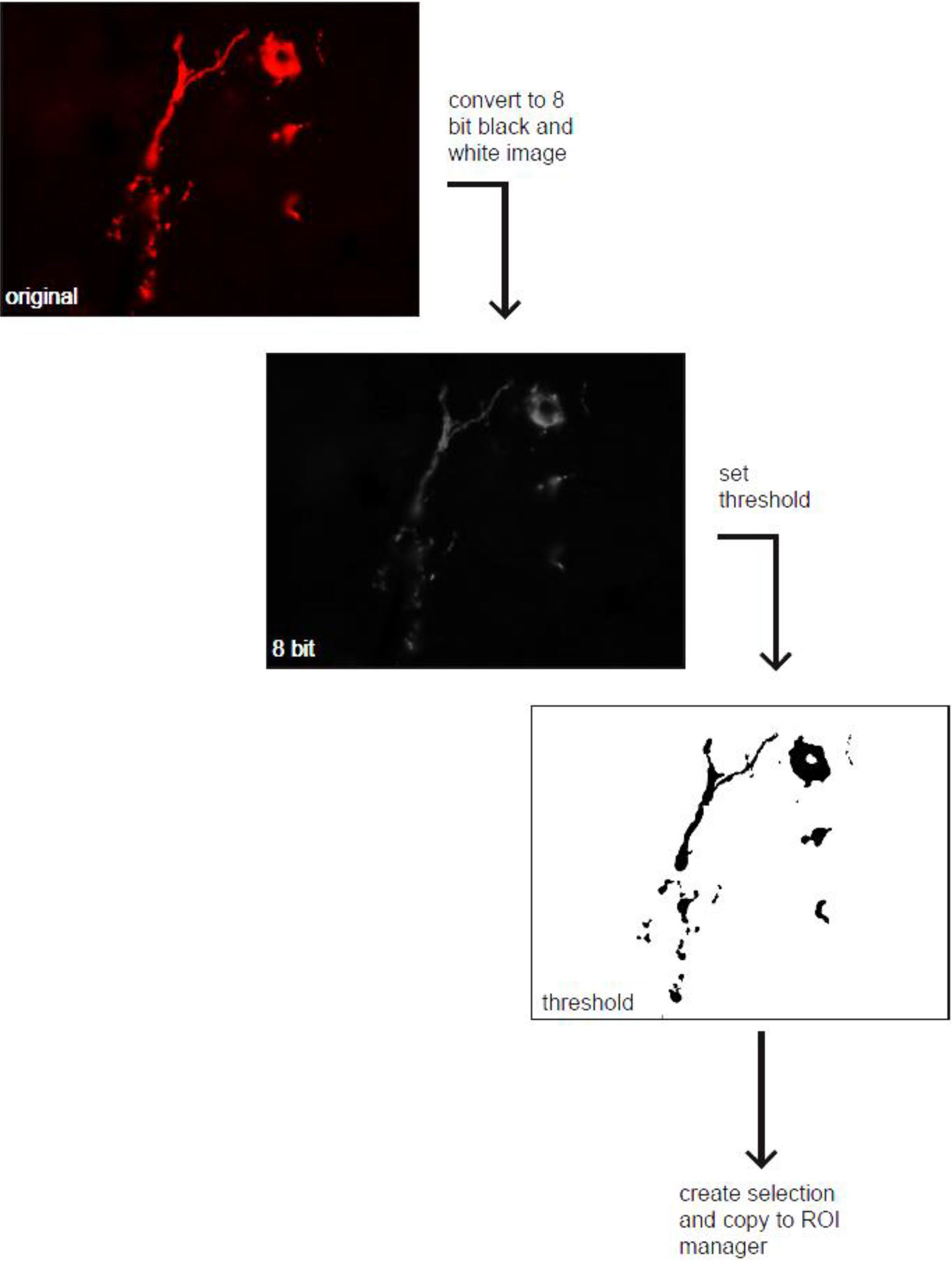
Schematic depiction of the mode of operation of the macro used to create ROIs of TH positive areas in the TH and peripherin double stained sections. First Images are converted to 8 bit black and white images then a threshold is set for brightness using the Otsu auto threshold tool of ImageJ. Finally a selection was created of the area that was above the threshold and added to the ROI manager tool of ImageJ. (Spleen tissue is shown in this example)

**Figure S2:**
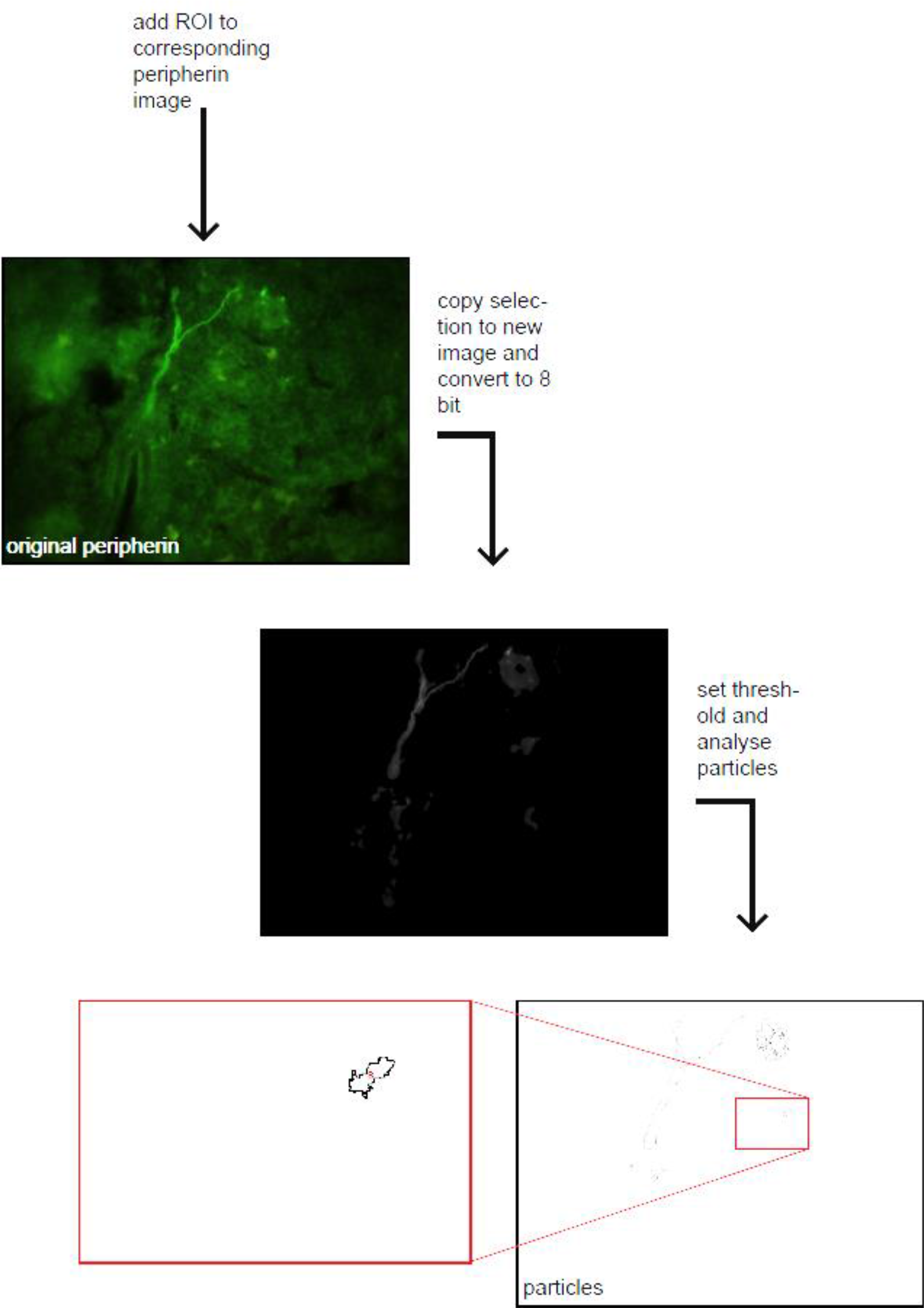
Schematic depiction of the mode of operation of the macro used to count the peripherin positive particles within the ROIs set by the previous macro. First the ROIs are placed on the peripherin image then a selection is created and copied to a new image. This image is converted to a bit and a threshold is set for brightness using the default auto threshold tool of ImageJ. The areas above the threshold are counted using the analyze particles tool of Image J. (Spleen tissue is shown in this example)

**Figure S3:**
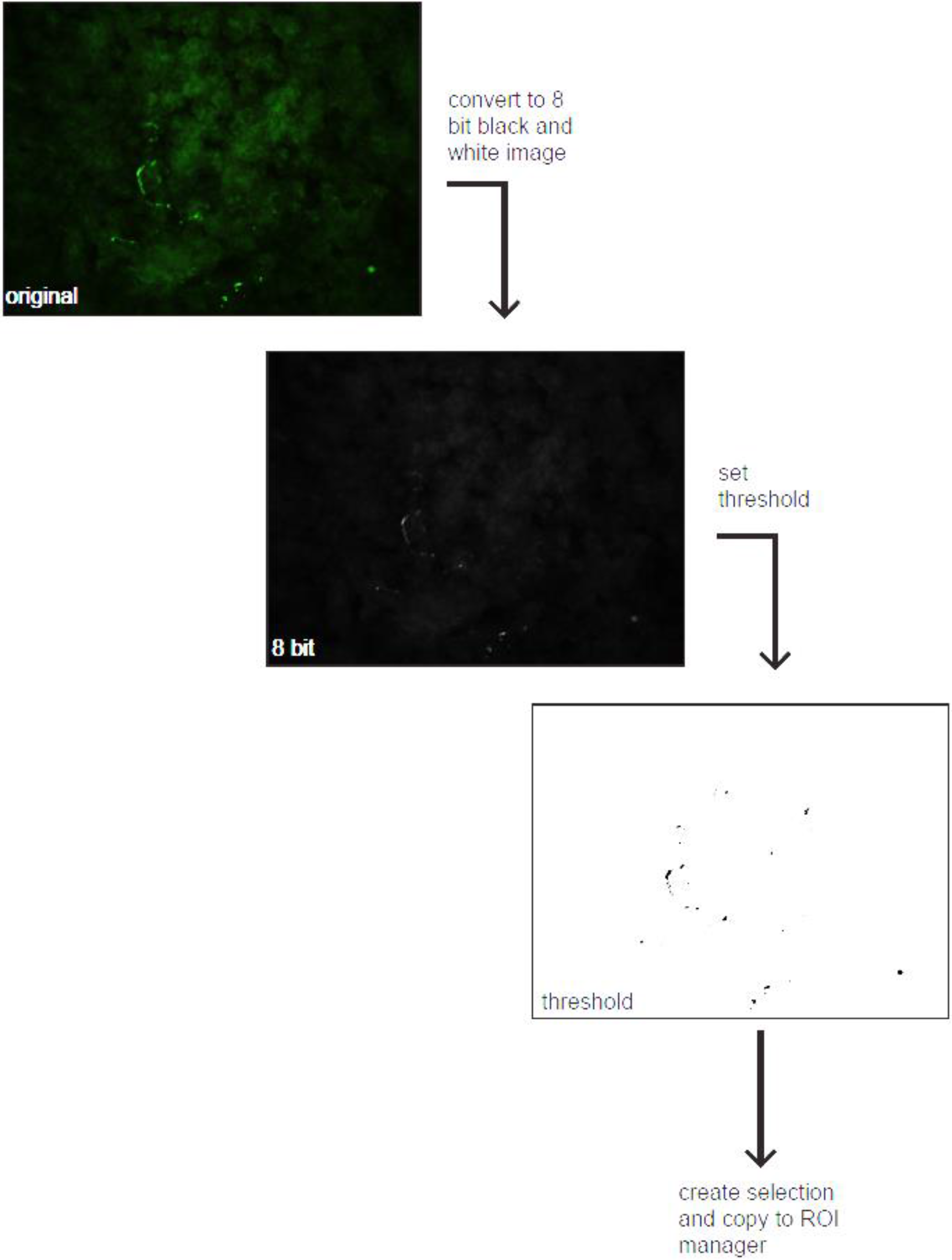
Schematic depiction of the mode of operation of the macro used to create ROIs of the peripherin positive areas in the TH and peripherin double stainings. First the images are converted to 8 bit black and white images and a threshold is set for brightness using the triangle auto threshold tool of ImageJ. Then a selection is created of the areas above the threshold and added to the ROI manager tool of ImageJ. (Spleen tissue is shown in this example)

**Figure S4:**
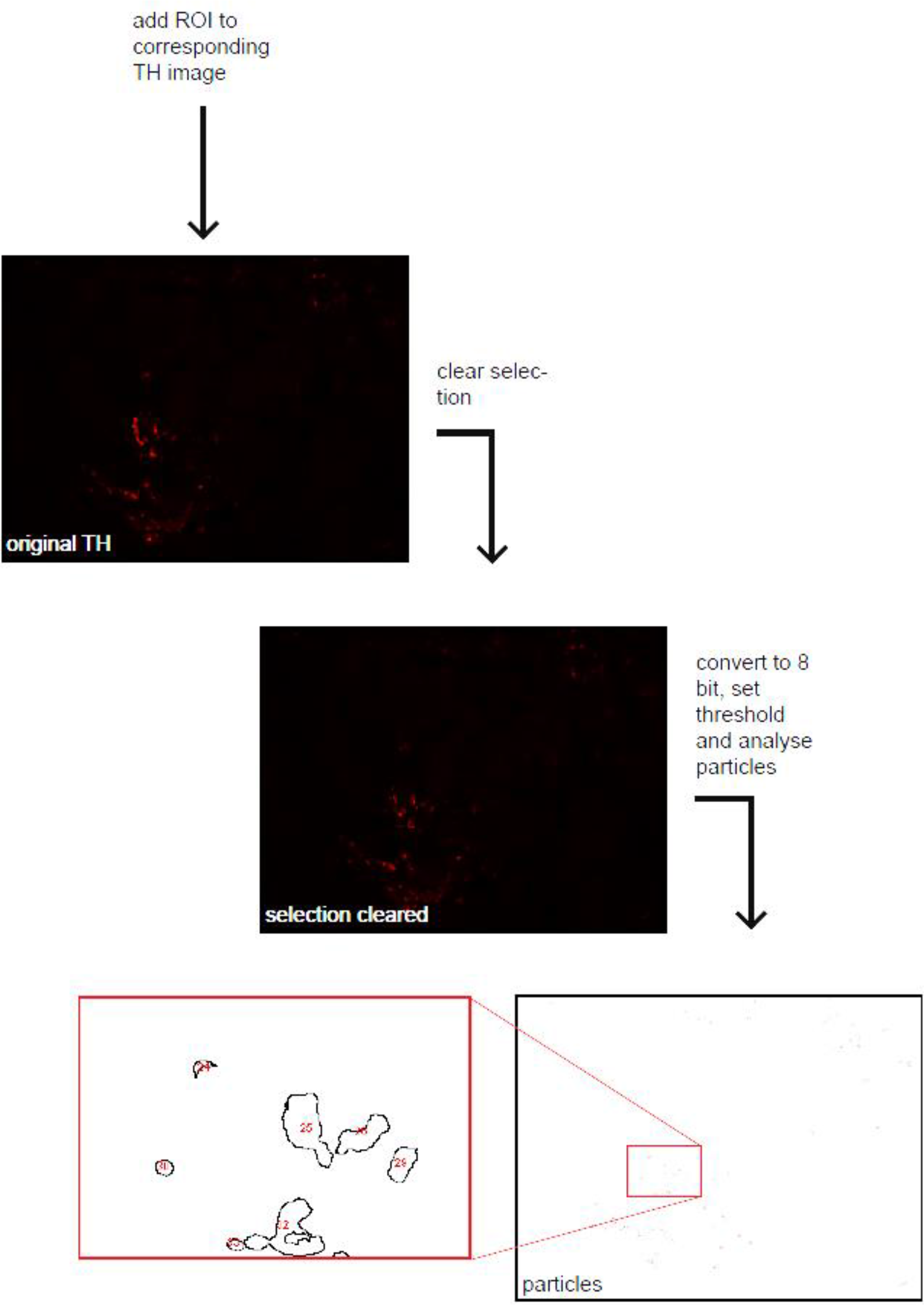
Schematic depiction of the mode of operation of the macro used to count TH positive particles after the ROIs of the peripherin positive areas from the previous macro have been removed. First the ROIs are placed over the TH images then a selection is made of these areas and cleared. The rest of the image is converted to an 8 bit black and white image and a threshold is set for brightness using the triangle auto threshold tool of ImageJ. The “analyze particles” tool is finally used to count particles above the threshold in brightness and above a defined size. (Spleen tissue is shown in this example)

**Figure S5:**
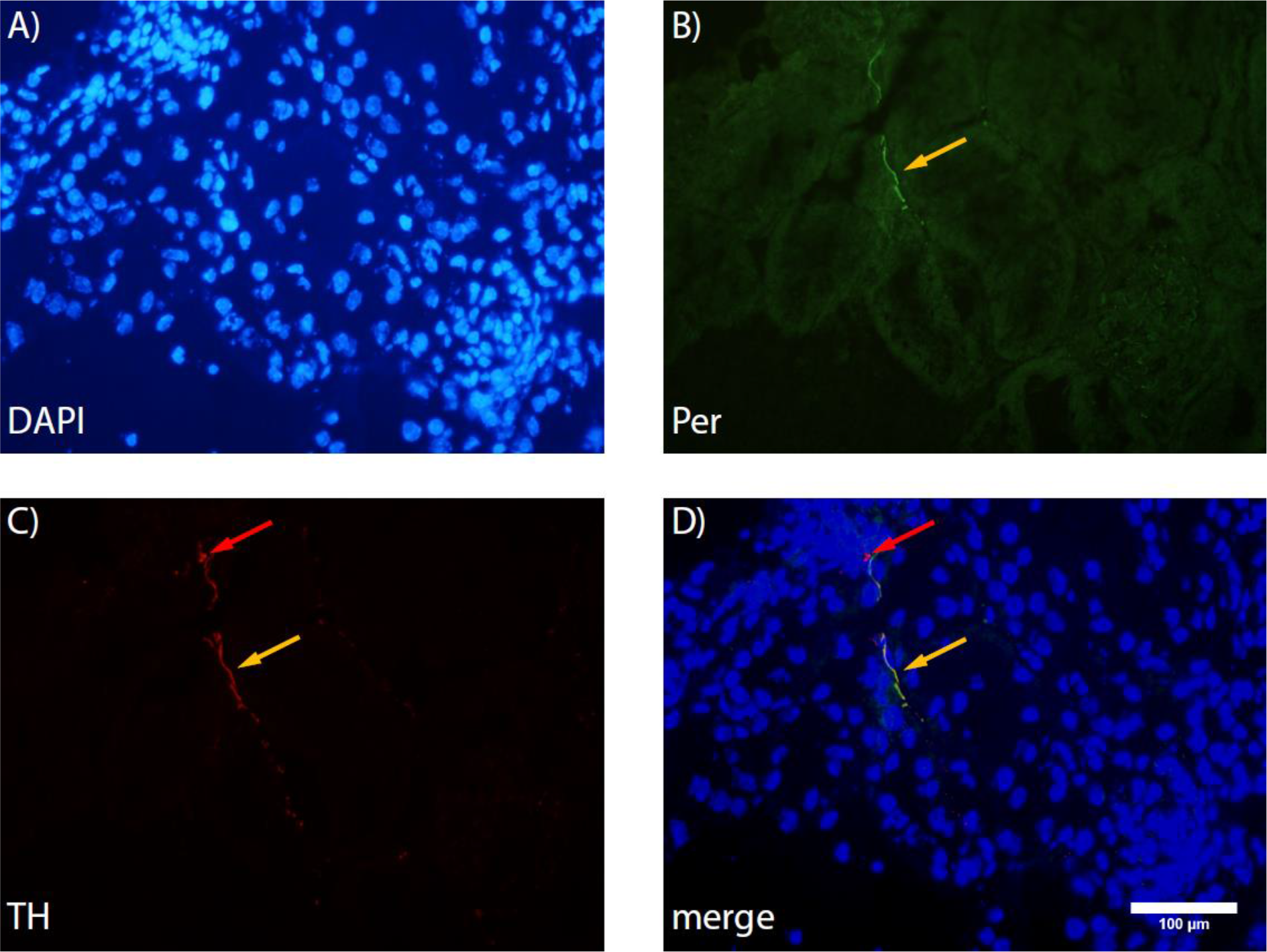
Exemplary image of TH – peripherin co – staining in an adrenal gland section. A) Nuclei are labeled with DAPI. B) Peripherin is labeled green (alexa fluor 488). C) TH is labeled red (alexa fuor 594). D) Merged image. Magnification is 400 fold. Red arrows indicate TH positive cells, yellow arrows indicate sympathetic fibers.

**Figure S6:**
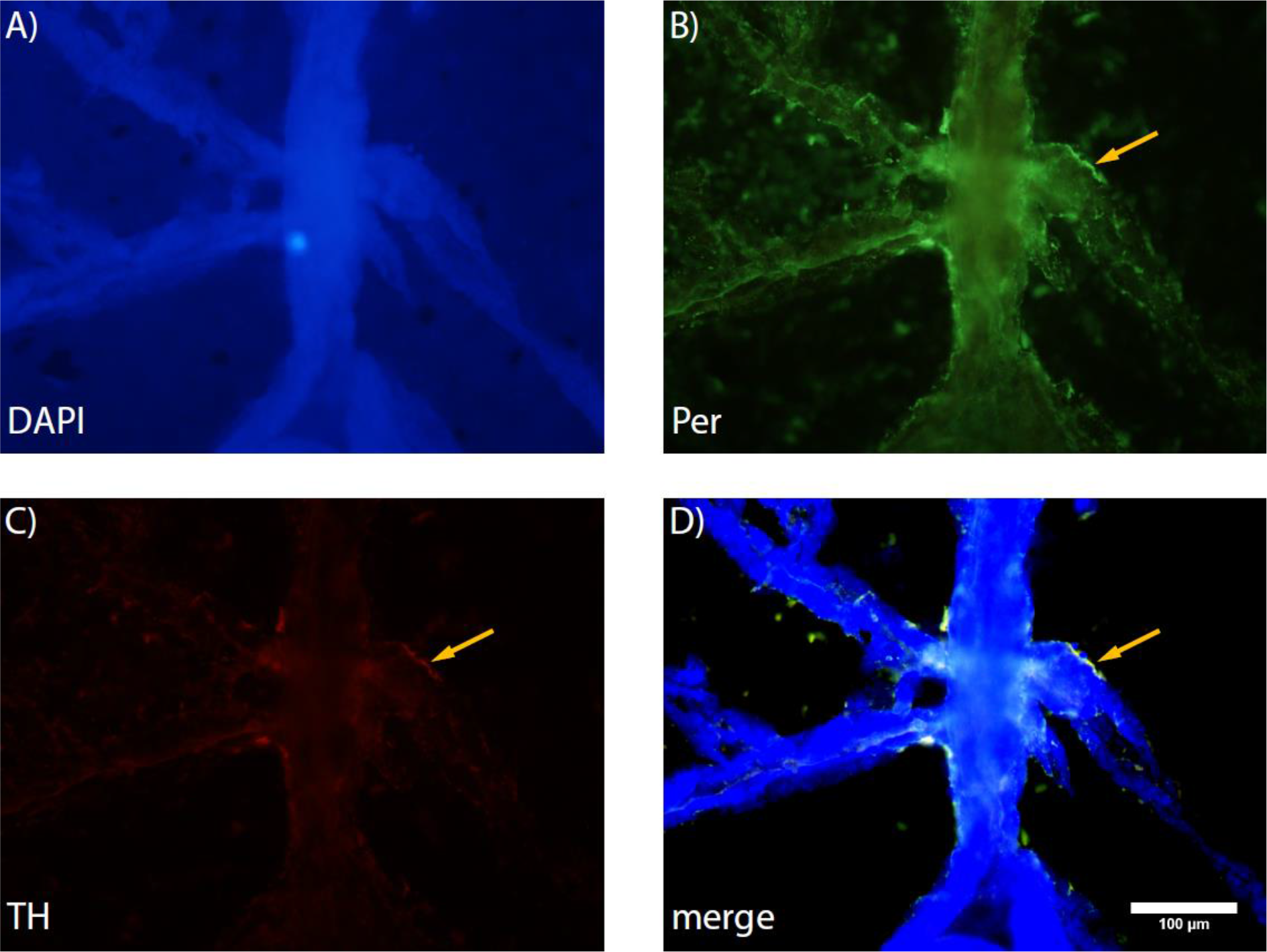
Exemplary image of TH – peripherin co – staining in a bone marrow (tibia) section. A) Nuclei are labeled with DAPI. B) Peripherin is labeled green (alexa fluor 488). C) TH is labeled red (alexa fuor 594). D) Merged image. Magnification is 400 fold. Yellow arrows indicate sympathetic fibers.

**Figure S7:**
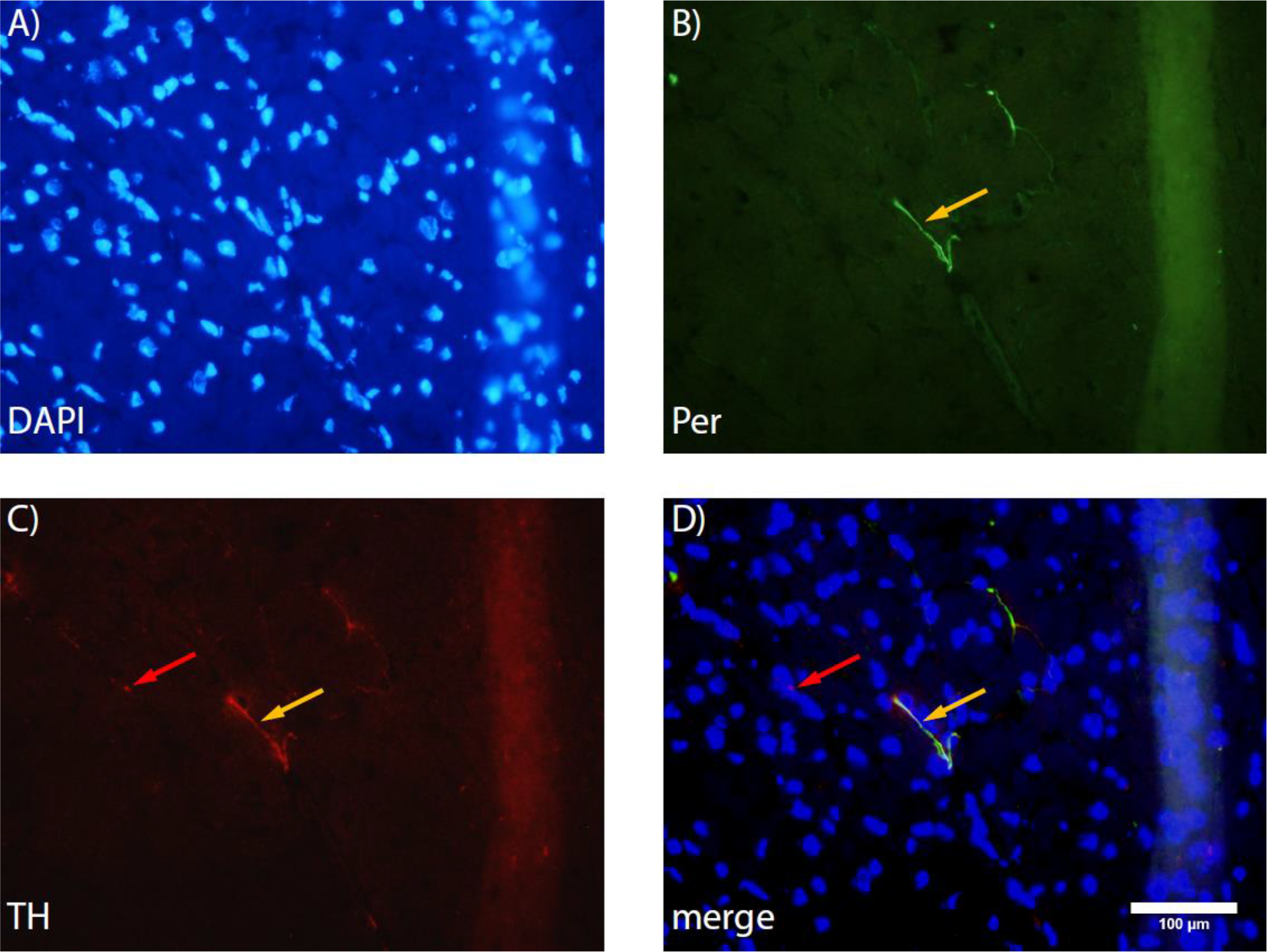
Exemplary image of TH – peripherin co – staining in a heart section. A) Nuclei are labeled with DAPI. B) Peripherin is labeled green (alexa fluor 488). C) TH is labeled red (alexa fuor 594). D) Merged image. Magnification is 400 fold. Red arrows indicate TH positive cells, yellow arrows indicate sympathetic fibers.

**Figure S8:**
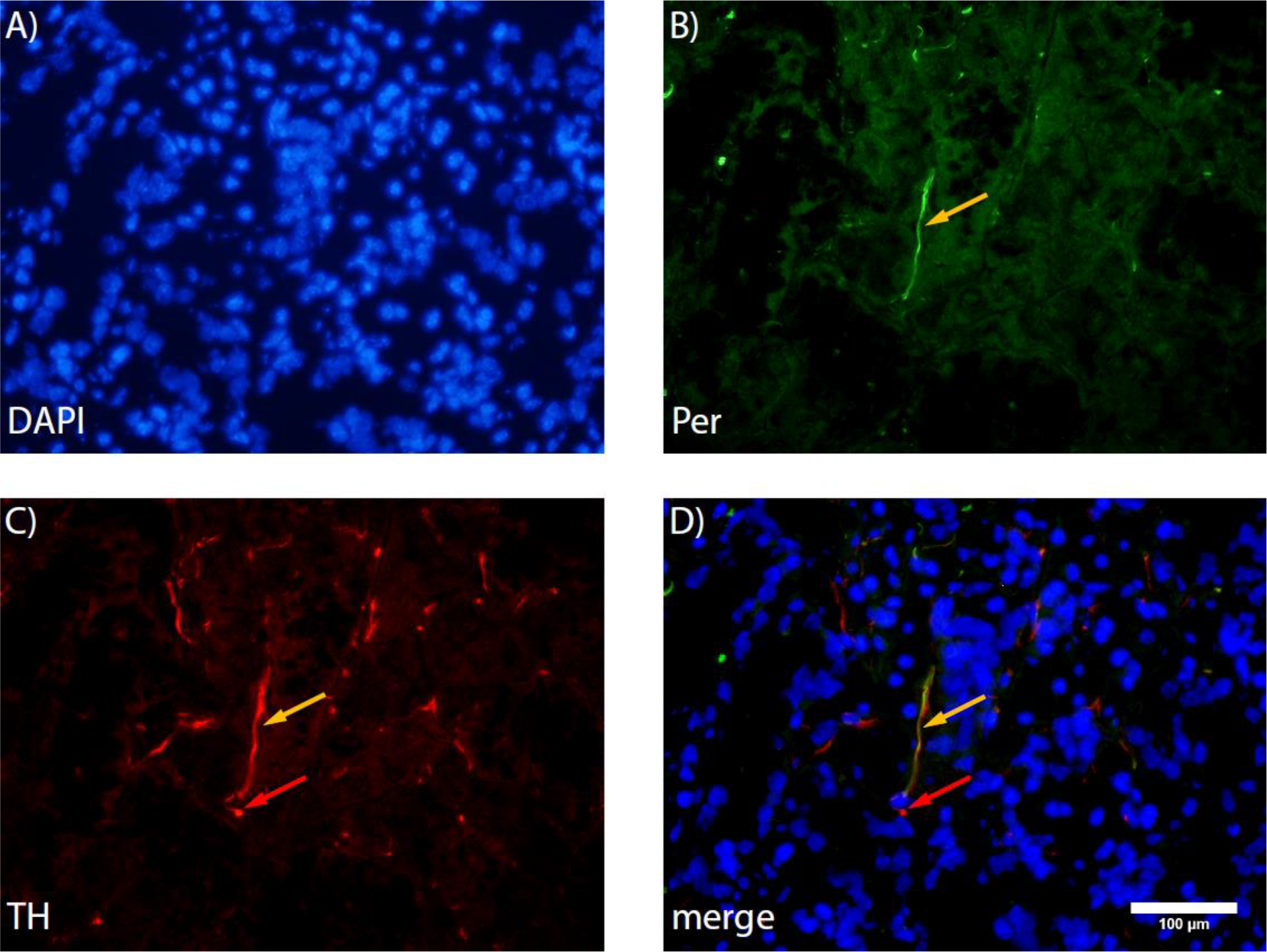
Exemplary image of TH – peripherin co – staining in a submandibular gland section. A) Nuclei are labeled with DAPI. B) Peripherin is labeled green (alexa fluor 488). C) TH is labeled red (alexa fuor 594). D) Merged image. Magnification is 400 fold. Red arrows indicate TH positive cells, yellow arrows indicate sympathetic fibers.

**Figure S9:**
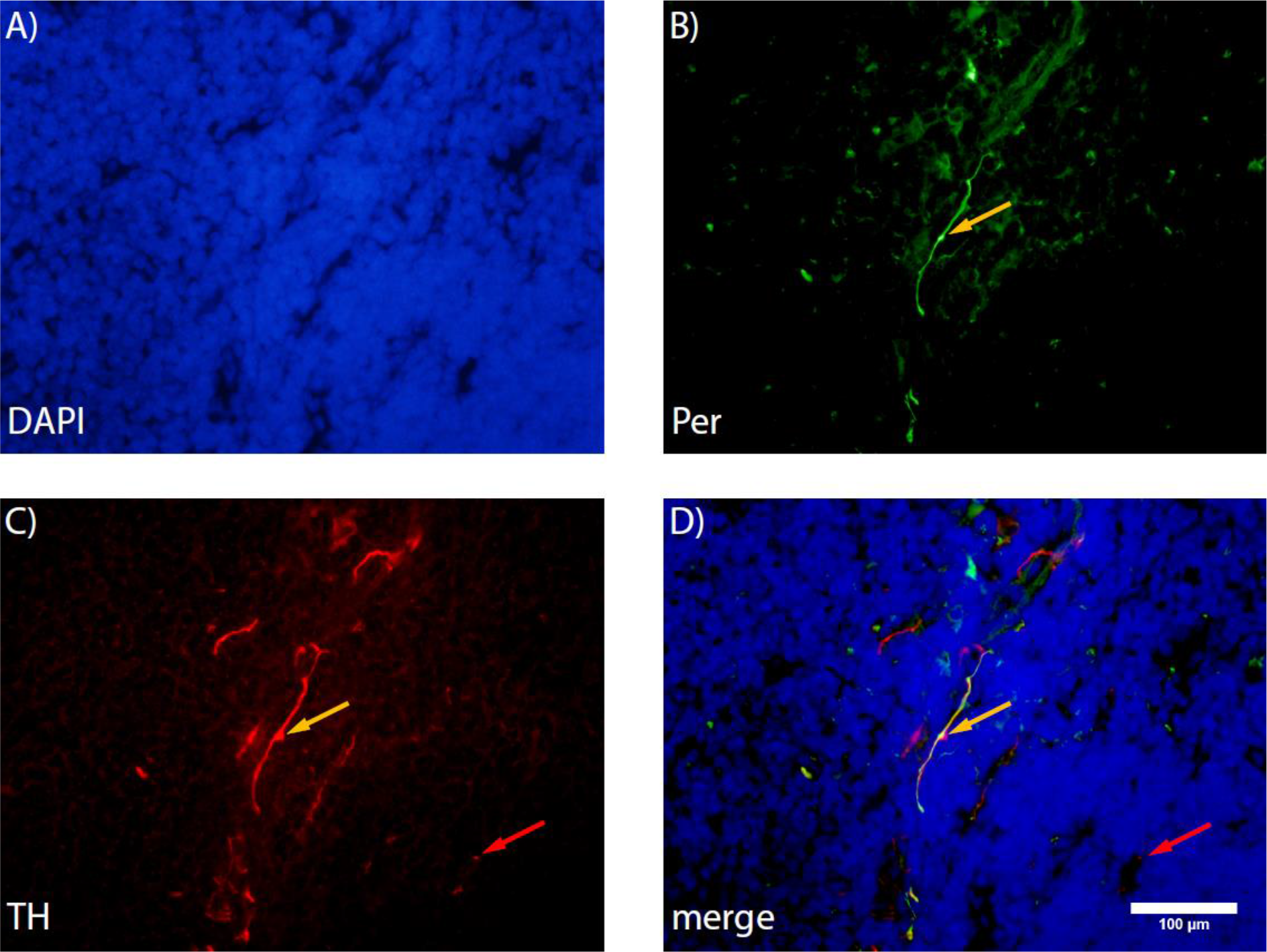
Exemplary image of TH – peripherin co – staining in a thymus section. A) Nuclei are labeled with DAPI. B) Peripherin is labeled green (alexa fluor 488). C) TH is labeled red (alexa fuor 594). D) Merged image. Magnification is 400 fold. Red arrows indicate TH positive cells, yellow arrows indicate sympathetic fibers.

